# Ranking of cell clusters in a single-cell RNA-sequencing analysis framework using prior knowledge

**DOI:** 10.1101/2023.10.02.560416

**Authors:** Anastasis Oulas, Kyriaki Savva, Nestoras Karathanasis, George M Spyrou

**Affiliations:** The Cyprus Institute of Neurology & Genetics, Bioinformatics Department, 6 International Airport Avenue, 2370 Nicosia, Cyprus, P.O. Box 23462, 1683, Nicosia, Cyprus

## Abstract

Prioritization or ranking of different cell types in a scRNA-Seq framework can be performed in a variety of ways, some of these include: i) obtaining an indication of the proportion of cell types between the different conditions under study, ii) counting the number of differentially expressed genes (DEGs) between cell types and conditions in the experiment or, iii) prioritizing cell types based on prior knowledge about the conditions under study (i.e., a specific disease). These methods have drawbacks and limitations thus novel methods for improving cell ranking are required. Here we present a novel methodology that exploits prior knowledge in combination with expert-user information to accentuate cell types from a scRNA-seq analysis that yield the most biologically meaningful results. Prior knowledge is incorporated in a standardized, structured manner, whereby a checklist is attained by querying MalaCards human disease database with a disease of interest. The checklist is comprised of pathways and drugs and, optionally, drug mode of actions (MOAs), associated with the disease. The user is prompted to “edit” this checklist by removing or adding terms (in the form of keywords) from the list of predefined terms. Our methodology has substantial advantages to more traditional cell ranking techniques and provides an informative complementary methodology that utilizes prior knowledge in a rapid and automated manner, that has previously not been attempted by other studies. The current methodology is also implemented as an R package entitled Single Cell Ranking Analysis Toolkit (scRANK) and is available for download and installation via GitHub (<u>https://#hub.com/aoulas/scRANK</u>)

**Author Summary:** Single-cell RNA Sequencing (scRNA-Seq) provides an additive resolution down to the cellular level that was previously not available from traditional “Bulk” RNA sequencing experiments. However, it is often difficult to prioritize the specific cell-types which are primarily responsible for the cause of a disease. This work presents a novel methodology that exploits prior knowledge for a disease in combination with expert-user information to accentuate cell types from a scRNA-seq analysis that are most closely related to the molecular mechanism of a disease of interest. Prior knowledge is incorporated in a standardized, structured manner, whereby a checklist is attained by querying a human disease database called MalaCards. This checklist comes in the form of pathways and drugs and, optionally, drug mode of actions (MOAs), associated with the disease. The expert-user is prompted to “edit” the prior knowledge checklist by removing or adding terms from the list of predefined terms.

This methodology has substantial advantages to more traditional cell prioritizing or ranking techniques and provides an informative complementary methodology that utilizes prior knowledge in a rapid and automated manner.

## Introduction

Single-cell RNA-sequencing (scRNA-seq) analysis generates clusters of cells that are grouped together due to their similar expression profiles. These clusters define specific cell types and provide scRNA-Seq data with increased resolution in comparison to bulk-RNA-Seq. This has apparent advantages when studying disease phenotypes because researchers can now narrow down the specific cell types that are affected by a given disease and perform more targeted, specialized analyses on them. The analysis of genes within specific cell types provides a very precise expression that is unique for the genes within the given cell type. However, it is often unclear which cell types may be more informative to investigate for a specific disease under study. It is common practice to perform a type of prioritization or ranking for the different cell types in a scRNA-Seq framework by: 1) obtaining an indication of the proportion of cells types between the different conditions under study (1), 2) counting the number of differentially expressed genes (DEGs) between cell types and conditions in the experiment (2) or, 3) prioritizing cell types based on prior knowledge about the disease (3). These methods have certain drawbacks and limitations such as: a) cell type proportion estimates from scRNA-seq data are variable. Statistical methods that can correctly account for different sources of variability are needed to confidently identify statistically significant shifts in cell type composition between experimental conditions (4). b) It has been shown that when using DEGs to prioritize cells, in both simulated and experimental datasets, the number of DEGs was strongly correlated with the number of cells per type, causing abundant cell types with modest transcriptional perturbations to be prioritized over rare but more strongly perturbed cell types (4). c) Using prior knowledge is limited to the selection of specific cell types known to be affected in a disease under study. This requires that the cell types are well documented and characterized from a clinical perspective. This is often not the case and most disease phenotypes are not so straightforward. Moreover, the specific type of cells that are mostly affected by the disease are not always so well documented.

Efforts have been made to improve prioritizing methods like the ones described above (4,5). However, no attempts have been made to improve the way prior knowledge is exploited in a scRNA-Seq framework. Currently, there is no structured and automated method of incorporating prior knowledge in scRNA-Seq analysis pipelines. Prior knowledge is often introduced in the analysis pipeline after manual inspection in order to select for the most informative cell types and consequently these cell types are used to perform downstream analyses (e.g., differential expression, pathway enrichment analysis, drug repurposing (DR)). The majority of studies do not use prior knowledge but instead adopt a discovery mode in their analysis pipeline (e.g., (1,2)). Results from these studies are difficult to validate as they do not coincide to previously documented information for the disease and it often takes the researchers thorough inspection to arrive to a concrete interpretation of the results.

In this study we present a method that exploits prior knowledge in combination with expert-user information to guide the choice of cell types from a scRNA-seq analysis that yield the most biologically meaningful results. Prior knowledge is incorporated in a standardized, structured manner, whereby a checklist is attained by querying MalaCards human disease database (6) with a disease of interest. The checklist is comprised of pathways and drugs and, optionally, drug mode of actions (MOAs), associated with the disease. The user is prompted to “edit” this checklist by removing or adding terms (in the form of keywords) from the list of predefined terms. The user may also define *de-novo*, a set of keywords that best suit a hypothesis the user is interested in investigating (hypothesis-driven approach). Once the checklist is finalized, a “mapping” step is performed. This is done initially against pathway enrichment results attained from analysing the scRNA-seq data. In addition, the user-selected checklist of drug names is mapped against transcriptomics-based drug repurposing (DR) results derived from the data. The analysis of scRNA-seq data has the capability to obtain pathways and repurposed drugs by comparing disease-control conditions for every cell type in the analysis. Our methodology then uses the user defined information to highlight the cell types that generate the results that best “map” to the predefined prior knowledge provided by the expert user. The methodology is fully automated and a ranking is generated for all cell types in the analysis allowing the user to pinpoint the specific cells that are most prominently affected by the disease under study in accordance to the provided prior knowledge (see Figure 1). The output of our methodology provides an automated validation of the result and further makes it easier for researchers to interpret their findings. In addition, the results provide greater credence to *de novo* information as it also backed-up by prior knowledge for the disease under study. Novel information is obtained in the form of predicted pathways that are enriched for each cell type, as well as previously unreported repurposed drugs that are obtained from the cell type-specific DEGs between disease-control conditions.

**Figure 1.**
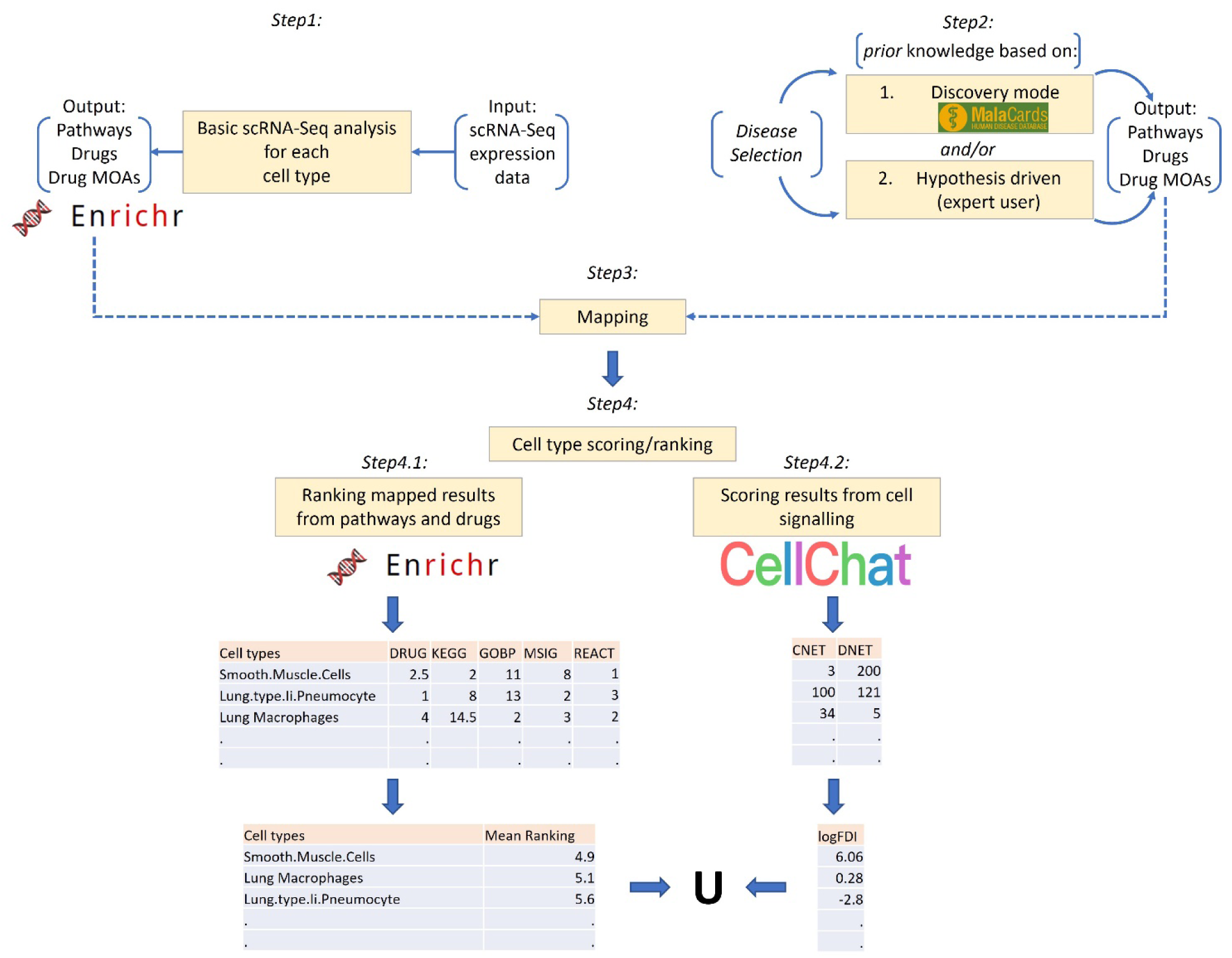
Flowchart of overall adopted methodology. *Step 1*: Basic scRNA-Seq analysis using SEURAT resulting in enriched pathways and repurposed drugs. *Step 2*: Defining prior knowledge. Two options are available: i) proceed with all the terms obtained from a check list of predefined terms provided by querying MalaCards with a disease of interest ii) to perform a hypothesis-driven approach whereby the user can provide specific keywords/terms associated with the hypothesis under investigation to perform a *de-novo* search across the supported databases. *Step 3*: Mapping step. Merges output from *Steps 1* and *2* and assesses how well prior knowledge “maps” to the results obtained from scRNA-Seq analysis. This is done firstly by mapping prior knowledge pathways against pathway enrichment results attained from analysing the scRNA-seq data. Secondly, prior knowledge drug names and/or drug mode of actions (MOAs) are mapped against drug repurposing results from analysing the scRNA-seq data using the CMAP database. *Step 3* is performed for all cell types in the analysis. *Step 4:* Scoring and ranking the cell types with respect to how well the data-driven output from pathway enrichment analysis and drug repurposing for individual cell types, “maps” to the predefined information provided by the expert user. *Step 4* is further split into 2 steps (4.1 and 4.2): *Step 4.1 –* Matching the position of the prior knowledge in the output (enriched pathways and repurposed drugs) of the scRNA-Seq analysis and then taking the Euclidian distance between the matched positions. *Step 4.2* – Ranking of cells using cell-cell communication networks generated using CellChat. Performing a comparison of the number of interaction (edges) between cell types (nodes) in the two different networks (control vs. disease) and ranking the cells by log fold difference in interactions (*LogFDI*) taking in consideration both positive and negative fold changes. Finally, the union between results is obtained (denoted by U above) taking into consideration the top 3 ranked cell types from *Steps 4.1* and *4.2*.

Some key points regarding our study are the following: 1) We developed a methodology that provides a link between scRNA-Seq data and public databases using prior knowledge. 2) This methodology allows for the efficient prioritization of cell types for a given disease of interest. 3) It facilitates the effective use of scRNA-Seq data for pathway enrichment analysis and DR and allows for the improved, targeted and highly-sensitive/specific prediction of pathways and repurposed drugs under a single-cell omics framework. 4) It provides a means of assessing the efficacy of enrichment analysis and DR performed using scRNA-Seq in comparison to bulk-RNA-Seq data.

## Materials and Methods

### Datasets

Three datasets were used in order to assess the performance of our methodology: COVID dataset 30 samples: Gene Expression Omnibus (GEO) accession GSE159812 (1). LAM dataset 8 samples: Gene Expression Omnibus (GEO) accession GSE135851 (3) Autism Dataset 41 samples: Sequence Read Archive, accession number PRJNA434002 (2).

### Basic Analysis Flow and Prior Knowledge Incorporation

The basic analysis flow of our methodology (see Figure 1 – *Step 1*) begins by performing scRNA-Seq analysis using SEURAT (ver. 4.3.0) (7). We utilized standardized analysis pipelines following best practices as described in the SEURAT web page (https://satijalab.org/seurat/index.html). The basic analysis includes, pre-processing, normalization, sample integration using anchors, dimensionality reduction via Principal Component Analysis (PCA), nearest-neighbor graph construction, clustering and detection of differentially expressed genes (DEGs). Custom-made scripts were used to perform in-script connections to enrichR via the corresponding R package (enrichR ver. 3.1) (8). The latter was done to attain annotation of cell-types based on marker gene analysis (if not supplied by the user) and also perform pathway enrichment analysis and DR. Next, the user is prompted to provide a set of “prior knowledge” information (see Figure 1 – *Step 2*). There are two options to consider: 1) Discovery mode: where the user proceeds with all the terms obtained from a checklist of predefined terms (default option). In this mode the checklist is obtained by querying MalaCards human disease database (6) with a disease of interest (e.g., Covid-19). The checklist is comprised of pathways and drugs, as well as drug MOAs (optional), associated with the disease. 2) Hypothesis-driven mode: where the user can provide specific keywords/terms associated with the hypothesis under investigation (i.e., to check for signs of inflammation in the different cell types, the user may provide targeted terms associated with inflammation, e.g., Cytokines). For the hypothesis-driven approach, the new keyword terms that are defined are used to perform de-novo querying across the different supported databases (see below). The search extracts the relevant database entries (if any) associated with the keywords. As it may be difficult for the user to provide specific drug names as keywords for this approach, our methodology supports the input of drug MOAs, which can be supplied instead. Our method then proceeds to search and extract the relevant drugs associated with these MOAs using information from the Drug Repurposing Hub Database (https://clue.io/repurposing). The Drug repurposing hub is a curated and annotated collection of FDA-approved drugs, clinical trial drugs, and pre-clinical tool compounds (9). This information is made available for our methodology as a text file.

The order of the prior knowledge information is important for downstream analytics. For the Discovery mode approach the order is defined by an enrichment score provided by MalaCards. For the Hypothesis-driven approach the user has to define the order of the keywords with most relevant/important keywords provided first.

### Mapping Prior Knowledge to Data-driven Results and Ranking of Cell Types

Once the prior knowledge information is finalized, the “mapping” step is performed (see Figure 1 – *Step 3*). This step merges *Steps 1* and *2* and whereby the output of *Step 1* is used as input for *Step 3* together with output from *Step 2*. This step assesses how well prior knowledge “maps” to the results obtained from scRNA-Seq analysis. This is done firstly by mapping prior knowledge pathways against pathway enrichment results attained from analysing the scRNA-seq data (data-driven analysis). Pathway enrichment is performed using selected DEGs and five pathway-related databases: (i) the Kyoto Encyclopedia of Genes and Genomes (KEGG) (10), (ii) Gene Ontology (GO) (11), (iii) the Molecular Signatures Database (MSIG) (12), (iv) WikiPathways (13) and Reactome (14). Secondly, prior knowledge on drug names and/or MOAs (depending on whether the *Discovery* or *Ηypothesis-driven* approach is undertaken) are mapped against DR results from the data-driven analysis using the CMAP database. Our methodology supports two options for *Step 3*. The default option generates different sets of results for every cell type in the dataset. There is also a “*Bulk-RNA*” option designed to simulate results generated from a bulk-RNASeq experiment. In this option results are generated by merging all different cell types together (per condition) and obtaining DEGs using these pooled cells across conditions.

Our methodology then uses the prior knowledge information to score/rank the cell types with respect to how well the data-driven output from pathway enrichment analysis and drug repurposing for individual cell types, “maps” to the predefined information provided by the expert user (see Figure 1 – *Step 4*). This is achieved by matching the position of the prior knowledge to the output of the scRNA-Seq analysis and then taking the Euclidian distance between the matched positions (see Figure 1 – *Step 4.1*). Cell-types that exhibit good mapping between prior knowledge and analysed results are characterized by small Euclidean distances (see Figure 2).

**Figure 2.**
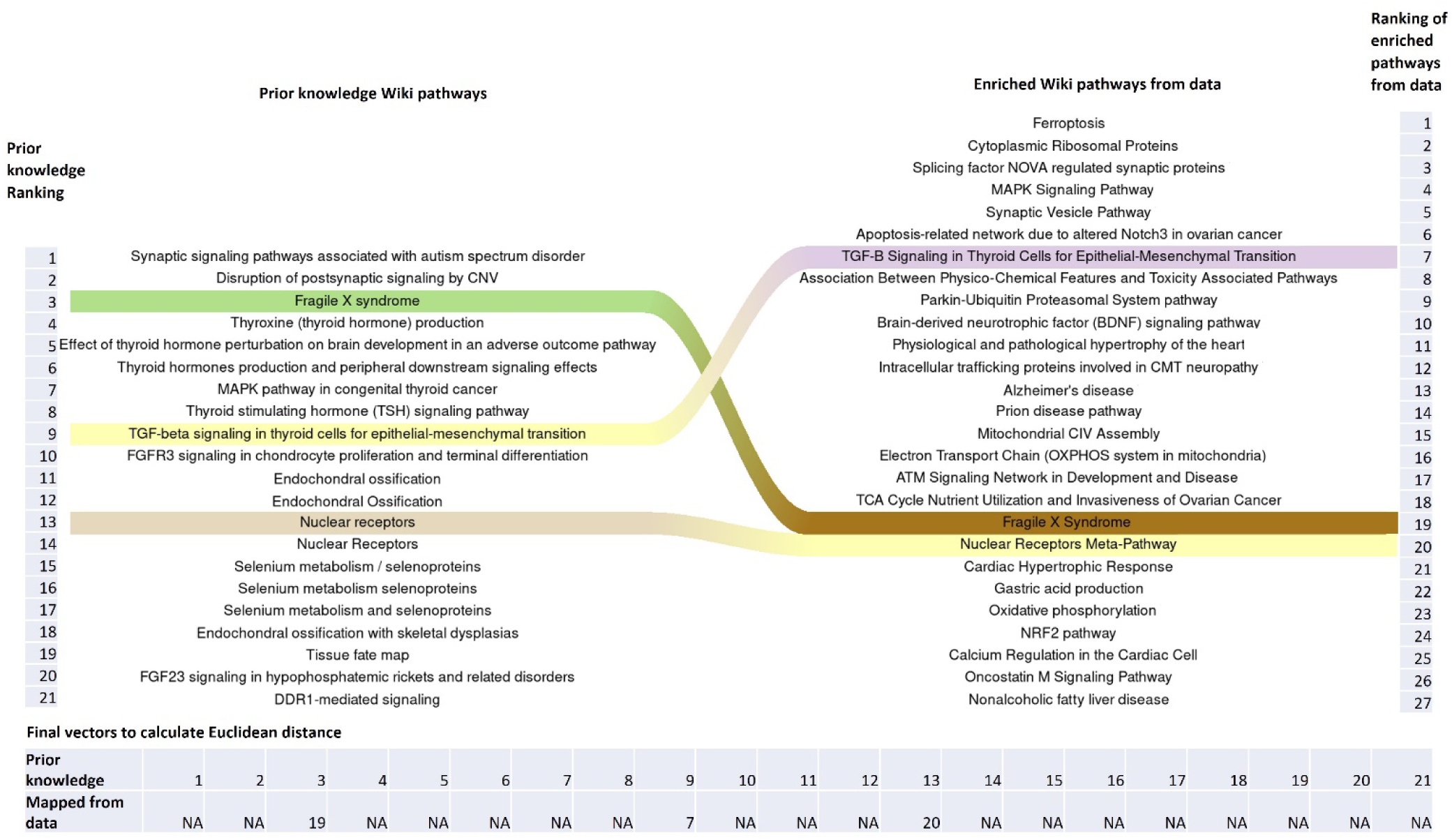
Riverplot showing an example of mapping of prior knowledge from MalaCards source wiki pathways to wiki pathways obtained from enrichment analysis on the scRNA-Seq data for specific cells type in a scRNA-Seq dataset. Mapping is performed by matching the position of the prior knowledge (left) to the output (enriched pathways) of the scRNA-Seq analysis (right) and then taking the *Euclidian* distance between the final vectors generated from the matched positions. Positions that are not matched/mapped at all receive a NA value.

In addition to this ranking step, cells are also ranked using cell-cell communication networks generated using CellChat (15) (see Figure 1 – *Step 4.2*). Global communications between cells are affected by disease. The assessment of these communications requires precise depiction of cell-cell signalling interactions and effective systems-level visualization. CellChat provides a database of interactions among ligands, receptors and their co-factors representing heteromeric molecular complexes. Furthermore, CellChat is capable of quantitatively analysing intercellular communication networks utilizing scRNA-seq data. It predicts major signalling inputs and outputs for cells and how those cells and signals coordinate towards the implementation of specific functions, using network analysis and pattern recognition approaches.

Capitalizing on the functionalities of CellChat, we first split scRNAseq data into control and disease samples and then generate two different types of cell-cell communication networks using CellChat. We then compare the number of interactions (edges) between cell types (nodes) in the two different networks (disease and control) and rank the cells by log fold difference in interactions (*LogFDI*). Ranking can be performed in either descending or ascending order as it is actually the absolute *LogFDI* value that determines the magnitude of the differences in interactions. The sign (+ve/-ve) merely dictates whether there is loss or gain of interactions with respect the control network. The *LogFDI* is obtained by calculating the log2 of the fold change of the interactions between cell types across disease and control conditions (see Equations 1, 2 and 3). The incentive here is to detect cell types whose signalling undergoes major alterations between disease and control conditions. These cells may be considered to be key effectors in the disease under study (see Figure 3). This provides a ranking which is based on topology information obtained from the CellChat networks. Finally, the different rankings are merged by obtaining the mean rank across all the rankings and a final ranking is assigned to each cell type of the analysis.

**Figure 3.**
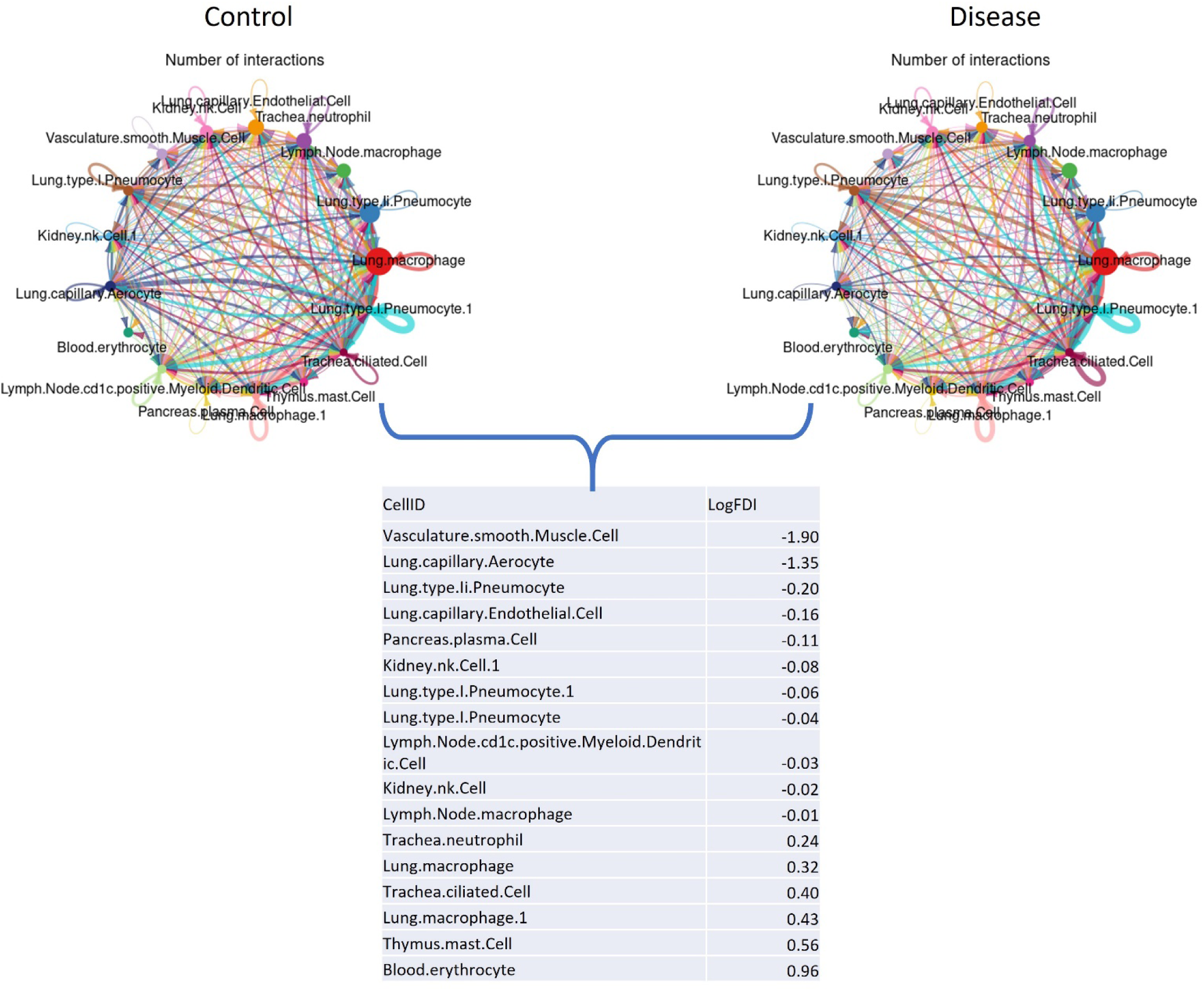
Example output from CellChat for a disease-control dataset. **A.** Data is split into two networks control and disease. **B.** Cell-types are ranked based on taking the log of the fold difference in their edge number interactions (*LogFDI*) between disease and control networks. Taking in consideration both positive and negative log fold changes. Node sizes are proportional to the number of cells in each cell group and edge widths with the number of interactions between nodes.

The log fold difference in interactions (*LogFDI*_*i*_) between disease and control networks for every cell type *i* is obtained as such:

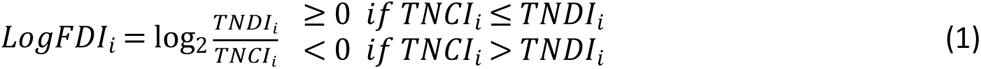

Where:

The normalized total number of control interactions for every cell type in the control network (*TNCI*_*i*_) is calculated as such:

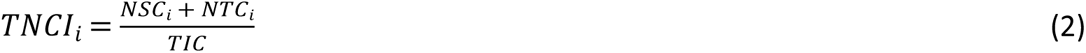

Where: *NSC*_*i*_ is the number of source-to-target interactions and *NTC*_*i*_ is the number of target-to-source interactions for each cell type *i* for the control network. *TIC* is the total number of interactions in the control network.

The normalized total number of disease interactions for every cell type in the disease network (*TNDI*_*i*_) is calculated as such:

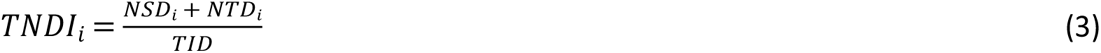

Where: *NSD*_*i*_ is the number of source-to-target interactions and *NTD*_*i*_ is the number of target-to-source interactions for each cell type *i* for the disease network. *TID* is the total number of interactions in the disease network.

### Bulk-RNA-Seq Simulations and other Cell Prioritization Methods

Each scRNA-Seq dataset was also evaluated using bulk-RNA-Seq simulations performed by merging all different cell types together and obtaining differentially expressed genes across conditions. The generation of this pseudo-bulk signature has been previously described (3) and is performed by differential expression between all disease cells and all control cells. This signature mimics the signature that would be obtained by the bulk-RNA-seq analysis. The Euclidian distances between the bulk-RNA-Seq data-driven results and the prior knowledge were treated as for the individual cell-types and a ranking obtained for the bulk-RNA-Seq.

In addition to prior knowledge, other methods are commonly used to prioritize cell types in a scRNA-Seq analysis framework. These are based on: 1) comparing the total number of DEGs between disease/control conditions for individual cell types and utilizing this as a means of ranking cell types and identifying the cells which are primarily affected by the disease under study (2), 2) ranking cell types by calculating the proportion of cell types across the different conditions in the analysis. For example, counting the number of different cell types between disease/control conditions can be an indication of the cellular differentiation/proliferation in disease vs. control conditions (1). Both of these approaches were assessed herein, for each dataset used as a case-study and the results were compared with our ranking methodology as well as the results reported by the authors who first published these datasets.

### R package implementation

The current methodology was implemented as an R package entitled Single Cell Ranking Analysis Toolkit (scRANK) and is available for download and installation via GitHub (<u>https://#hub.com/aoulas/scRANK</u>). The basic analysis function supports data in multiple output formats obtained from Cell Ranger (16), namely: txt, csv, as well as H5 format. The methodology also supports pre-processed SEURAT R objects, so the user has multiple options by which to upload data into our pipeline. Moreover, if annotated datasets exist, scRANK supports the upload of meta.txt files containing SEURAT metadata for the corresponding dataset.

## Results

Public datasets were downloaded in order to evaluate different case studies for validating our methodology. Every case study was evaluated by a) Using the default option (Discovery Mode) whereby information derived from MalaCards was used as prior knowledge input and b) Using the Hypothesis-driven Mode whereby an expert user can manually input keywords as prior knowledge in order to extract information from the relevant databases, thus, allowing for the investigation of specific hypotheses of interest. In order to evaluate the ranking of cell types by our methodology in comparison to bulk-RNA-Seq data, we further performed bulk-RNA-Seq simulations. This was done by merging all different cell types and obtaining differential expressed genes across conditions for each of the datasets used as a case study. The Euclidian distances between the bulk-RNA-Seq data-driven results and the prior knowledge were calculated and compared to the Euclidian distances attained from the individual cell-types.

### Case study 1 – LAM disease

#### Discovery Mode Approach

Lymphangioleiomyomatosis (LAM) is a lung disease caused by the abnormal growth of smooth muscle cells, especially in the lungs and lymphatic system. This abnormal growth leads to the formation of holes or cysts in the lung. Causative mutations for LAM disease are known to be present in TSC1 or TSC2 genes. These cause the hyperactivation of the mTOR complex 1 (mTORC1). Under normal circumstances, mTOR is a major regulator of cell growth and division. The target of rapamycin (TOR) signal-transduction pathway is an important mechanism by which eucaryotic cells adjust their protein biosynthetic capacity to nutrient availability. Both in yeast and in mammals, the TOR pathway regulates the synthesis of ribosomal components, including transcription and processing of pre-rRNA, expression of ribosomal proteins and the synthesis of 5S rRNA. However, in LAM cells, abnormally activated mTOR sends signals that encourage cells to grow uncontrollably. Activators of mTOR involve PI3K, phosphatidylinositol-dependent kinase-1 (PDK1), and Akt. Dysregulated mTOR leads to the hyperactivation of multiple pathways, including the MAPK signalling cascade and hence to downstream growth-related perturbations. The mTORC1 inhibitor Sirolimus is the only FDA-approved drug to treat LAM. Novel therapies for LAM are urgently needed as the disease recurs with discontinuation of the treatment and some patients are insensitive to the drug. When studying LAM it may be more informative to look at the expression profiles of genes as they are expressed specifically in smooth muscle cells, as these are the primary affected cells for this type of disease (17–19). This was the approach adopted by the authors that first published this dataset.

We further assessed the LAM dataset using our methodology in order to 1) Incorporate prior knowledge on LAM disease and perform ranking of the cell types identified in the SEURAT analysis. 2) Obtain additional repurposed drugs that can potentially be used in LAM disease.

The prior knowledge which was included as a list of terms for the LAM disease dataset was obtained by querying MalaCards for the disease “Lymphangioleiomyomatosis”.

The full list of pathways, drugs and MOAs (see supplementary Tables S1-5) obtained from the search was then used as input for our methodology to perform cell-ranking. These keywords best characterize the pathways, biological processes, as well as the drugs and drug MOAs, that are known to be implicated in LAM disease (according to MalaCards).

Results from the rankings of cell types using pathways and drug information show that our results are in agreement with the results reported by the authors who first published this dataset (3) as well as the clinical phenotype of the disease (17–19). Smooth muscle cells attained the highest ranking with respect to the rest of the cell types in the analysis (see Figure 4A). Results of the enrichment analysis and DR for some of the LAM top ranked cells are shown as supplementary Figure S1-3.

**Figure 4.**
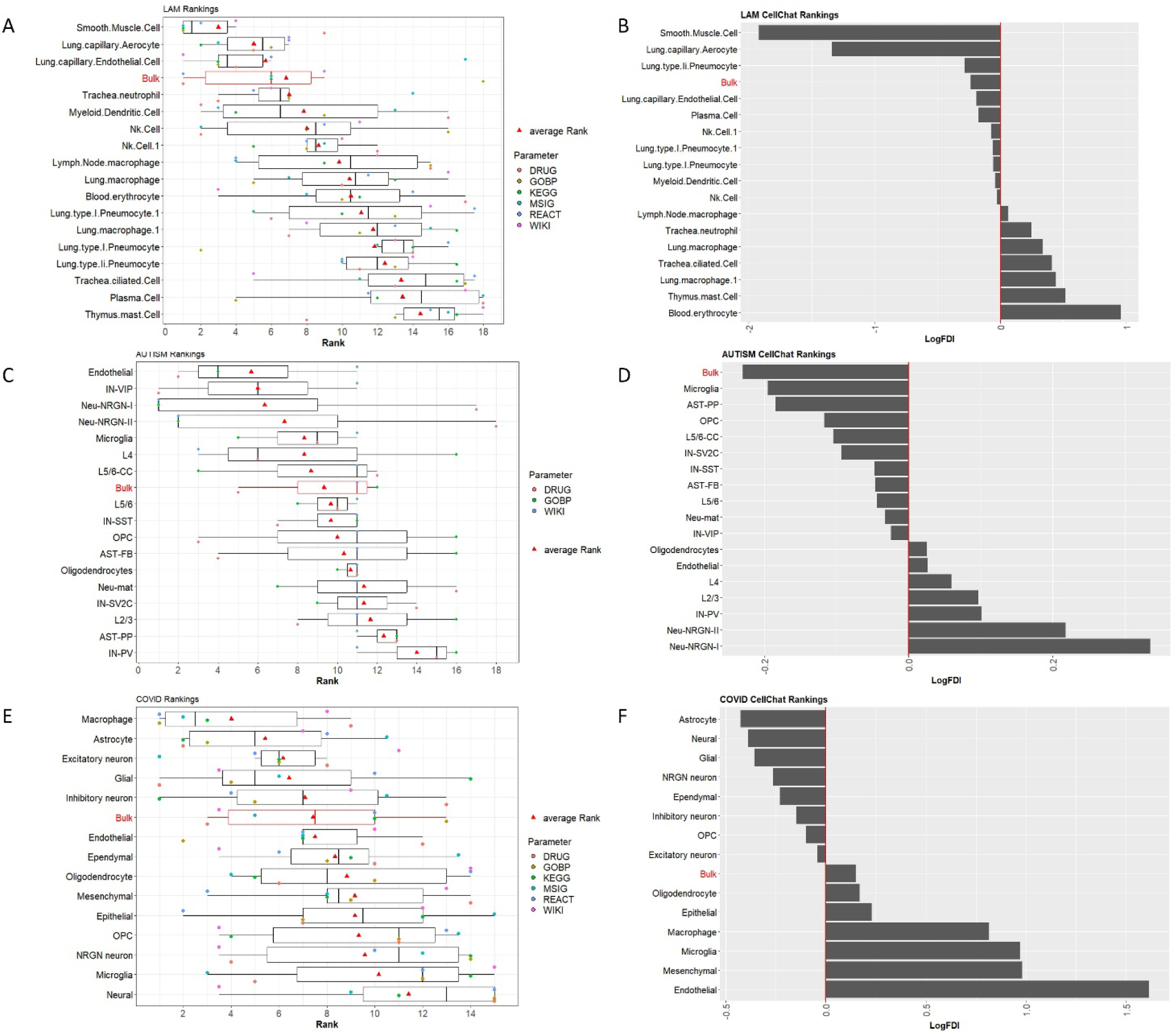
Discovery mode approach rankings across all 3 datasets. **A**. LAM cell-rankings based on drugs (DRUG) and pathway databases (KEGG, GOBP, MSIG, WIKI, REACT). The boxplot shows the individual cell rankings obtained by our methodology. The order of the cells is defined by the average rankings across all six parameters used to generate the final ranking for the annotated cell types in the analysis. **B.** CellChat rankings of LAM dataset cell-types based of *LogFDI*. **C**. ASD cell-rankings based on drugs and pathway databases (WIKI and GOBP – Note that KEGG, MSIG and REACT were not successfully mapped with prior knowledge for this case study). The boxplot shows the individual ranking obtained by our methodology. The order of the cells types is defined by the average rankings across the three parameters that attained mapping information in the analysis. **D.** CellChat ranking of autism dataset cell-types based on *LogFDI*. **E.** COVID cell-rankings based on drugs (DRUG) and pathway databases (KEGG, GOBP, MSIG, WIKI, REACT). The boxplot shows the individual ranking obtained by our methodology. The order of the cells is defined by the average rankings across all six parameters used to generate the final ranking for the annotated cell types in the analysis. **F.** CellChat rankings of COVID-19 dataset cell-types based on *LogFDI*. Results from the bulk RNA-Seq simulation are also shown (boxplots highlighted in red).

Interesting results were obtained from comparing number of signalling interactions using the cell-cell communication tool CellChat. Results showed that smooth muscle cells sustained the most differences between donor and LAM conditions, with a markable >3-fold decrease in the normalized number of interactions in the LAM condition compared to the donor (see Figure 4B). It appears that a lot of cell-cell communications pathways may be hindered in these cells under LAM conditions, resulting in loss of cell-cell interactions and denoted as decrease in the number of edge connections in the CellChat LAM network. Smooth muscle cells are the major effectors of LAM disease; as reported in clinical conditions (17–19), hence, perhaps it is apparent for these cell-types to sustain the greatest loss in connectivity when compared to other cell-types in the analysis. Results from the LAM bulk RNA-Seq simulation are also shown in the box-plots (see Figure 4A). In this case, we observe a relatively high ranking, albeit lower than the most informative cell –types for the disease under study.

#### Hypothesis-Driven Approach

The prior knowledge is included as keywords for the hypothesis-driven approach in order to query: 1) the five pathway databases (KEGG, GOBP and MSIG Hallmark, WIKI and REACT). 2) The drug repurposing databases (CMAP).

The prior knowledge keywords for LAM disease used for searching the five pathway databases were:

**Keywords KEGG**: MTOR, PI3K, MAPK, apoptosis, NF-k and TNF.

**Keywords GO**: MTOR, PI3K, MAPK, apoptosis, NF-k and TNF.

**Keywords MSIG**: MTOR, PI3K, MAPK, apoptosis, NF-k and TNF.

**Keywords REACT**: MTOR signalling, PI3K, MAPK, apoptosis, NF-k and TNF.

**Keywords WIKI**: MTOR, PI3K, MAPK, apoptosis, NF-k,TNF.

As mentioned in the Material and Methods (see above) for the hypothesis-driven approach, the keywords for searching drugs are supplied in the form of drug MOAs and then mapped to the relevant MOAs extracted from CMAP. The keywords used for this case study where: **Keywords (MoAs):** CDK inhibitor, MTOR inhibitor, MEK inhibitor.

The full list of pathways and drug MOAs (data not shown) obtained from the search were then used as input for our methodology to perform mapping and consequently cell-ranking. These keywords best characterize the pathways, biological processes, as well as the drug MOAs, that are known to be implicated in LAM disease (in accordance to LAM bibliography (17–19)).

Results from the rankings of cell types via the hypothesis-driven approach are in agreement with the results obtained by the Discovery Mode approach and are also in alignment with the results reported by the authors who first published this dataset (3), with lung capillary aerocytes together with smooth muscle cells attaining the highest ranking with respect to the rest of the cell types in the analysis (see Figure 5A).

**Figure 5.**
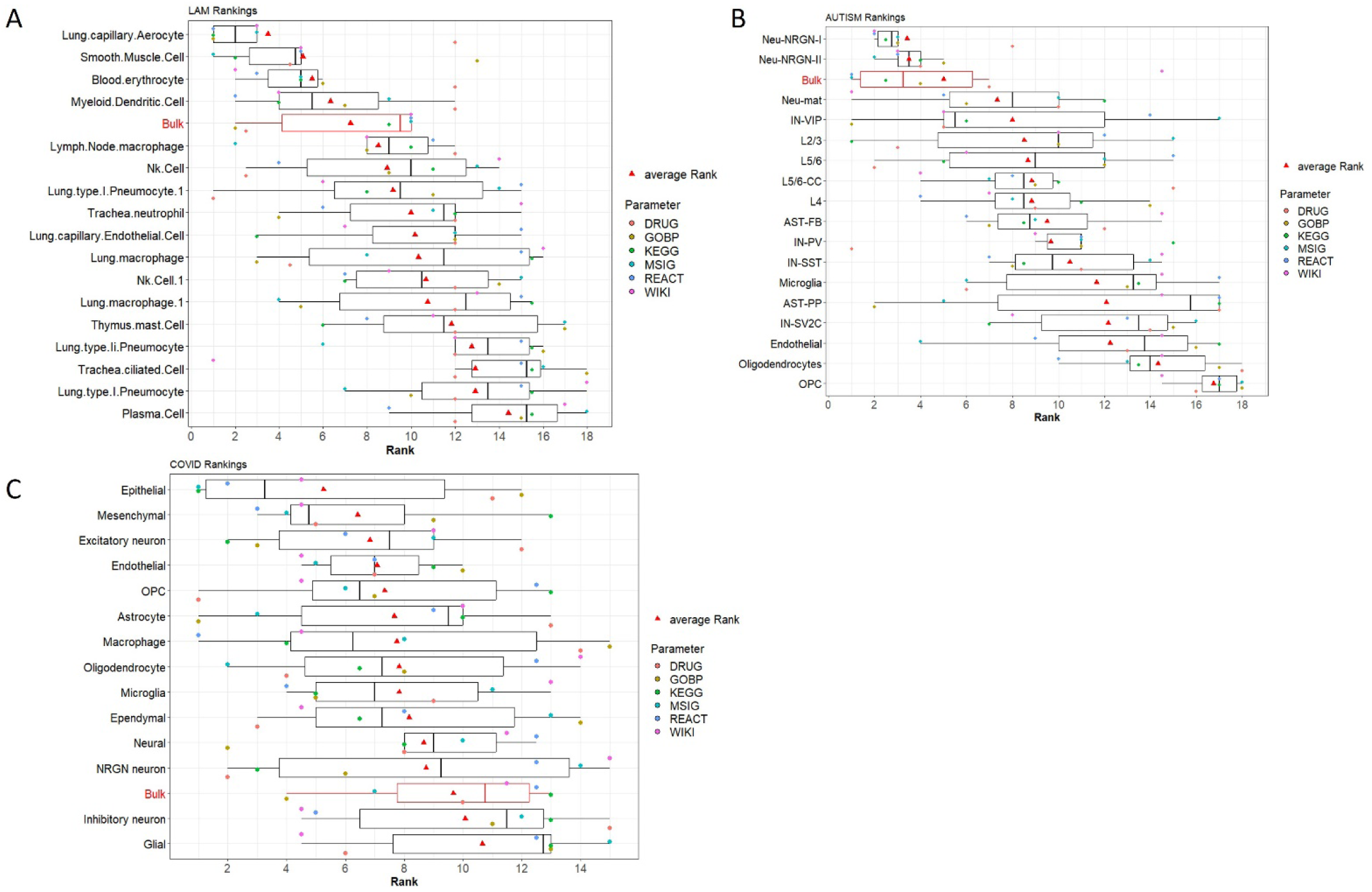
Hypothesis-driven rankings across all 3 datasets. **A**. LAM cell-rankings based on drugs (DRUG) and pathway databases (KEGG, GOBP, MSIG, WIKI, REACT). The boxplot shows the individual cell rankings obtained by our methodology. The order of the cells is defined by the average rankings across all six parameters used to generate the final ranking for the annotated cell types in the analysis **B**. ASD cell-rankings based on drugs and pathway databases (KEGG, GOBP, MSIG). The boxplot shows the individual ranking obtained by our methodology. The order of the cells is defined by the average rankings across the three parameters that attained informative information used to generate the final ranking for the annotated cell types in the analysis. **C.** COVID-19 cell-rankings based on drugs (DRUG) and pathway databases (KEGG, GOBP, MSIG, WIKI, REACT). The boxplot shows the individual ranking obtained by our methodology. The order of the cells is defined by the average rankings across all six parameters used to generate the final ranking for the annotated cell types in the analysis. Results from the bulk RNA-Seq simulation are also shown (boxplots highlighted in red).

### Case study 2 – Autistic Spectrum Disorder (ASD)

#### Discovery Mode Approach

Clinical classification of patients with autistic spectrum disorder (ASD) is based on current WHO criteria and provides a valuable but simplified depiction of the true nature of the disorder. A recent study (2) has shown that single-nucleus RNA sequencing of cortical tissue from patients with autism can be used to significantly enhance the resolution provided by bulk gene expression studies. Overall, the scRNA-Seq study shows that changes in the neocortex of autism patients converge on common genes and pathways. The authors report that synaptic signalling of upper-layer excitatory neurons (L2/3), vasoactive intestinal polypeptide (IN-VIP)–expressing interneurons and the molecular state of microglia are preferentially affected in autism. Moreover, results show that dysregulation of specific groups of genes in L5/6 cortico-cortical projection neurons correlates with clinical severity of autism.

As with the previous case study, prior knowledge is included as keywords for ASD as obtained from querying MalaCards with the diseases “Autism Spectrum Disorder” as well as “Autism”. As these terms are synonymous both keywords were used to query MalaCards

We obtained a list of terms that best characterize the pathways, biological processes, as well as drugs that are known to be implicated in ASD disease (with respect to MalaCards). The full list of pathways and drugs (see supplementary Tables S6-10) obtained from the search was then used as input for our methodology to perform cell-ranking.

Similarly to the LAM dataset analysis, the results from the rankings of cell types using pathways and drug information show that our results are in alignment with the results reported by the authors who first published the ASD dataset (2). The authors report that top differentially expressed neuronal genes were down-regulated for ASD specifically in L2/3 excitatory neurons and vasoactive intestinal polypeptide (IN-VIP)–expressing interneurons. The top genes differentially expressed in non-neuronal cell types were up-regulated for ASD in protoplasmic astrocytes and microglia. They also report cell types that are recurrently affected across multiple patients as upper (L2/3) and deep layer (L5/6-CC) cortico-cortical projection neurons. This is also seen in our results whereby, IN-VIP cells attained high ranking with respect to the rest of the cell types in the analysis (see Figure 4C). We did not achieve high ranking for the L2/3 cell, reported in the ASD study and perhaps this can be attributed to the lack of mapped information for the pathways extracted from the MalaCards database. Enrichment analysis and DR results for some of the ASD top ranked cells are shown as supplementary Figure S4-7. Results from the ASD bulk RNA-Seq simulation are also shown in the box-plots (see Figure 4C) attaining a middling ranking with respect to the individual cell-types in the analysis.

Results from comparing the normalized number of signalling interactions between ASD and control networks showed minor differences between disease and control networks. Pseudo bulk RNA analysis generated the highest differences, while from the cell types; Microglia displayed some differences between control and disease conditions showing a 1.14-fold decrease in the normalized number of interactions in the ASD compared to the control network (see Figure 4D). The authors report that the molecular state of microglia is preferentially affected in ASD and they are also tightly correlated with clinical severity of the disorder. As ASD is also linked to inflammation (20–22), it is not surprising to observe such changes in the primary innate immune cells of the brain. Moreover, Neu-NRGN-I and II cells attained notable decrease in fold change of normalized interactions between disease control networks with respect to the rest of the cell types in the analysis (see Figure 4D). These cells were not previously reported to be implicated in autism by the authors of this dataset in their original study. These are Neurogranin (Ng) expressing neurons, which is encoded by the schizophrenia risk gene *NRGN*. Ng is critical for encoding contextual memory and regulating developmental plasticity in the primary visual cortex during the critical period. The overall impact of Ng on the neuronal signalling and inflammation that regulates synaptic plasticity is unknown (23). This can perhaps be attributed to the lack of available molecular pathways and drugs currently known to be implicated in ASD.

#### Hypothesis-Driven Approach

As with the previous case study, prior knowledge is included as keywords (pathways and drug MOAS) and our methodology was applied. The prior knowledge keywords for ASD in order to search pathway databases (KEGG, GOBP, MSIG Hallmark, WIKI and REACT) were:

**Keywords KEGG**: Pathways of neurodegeneration, Prion, Alzheimer, Parkinson, ALS, axon.

**Keywords GO**: axon, synapse, neurotransmitter, neuron, PI3K, MAPK, apoptosis, NF-k, TNF, JAK, STAT, Cytokine, Inflammation, Th17, Th1, IL-.

**Keywords MSIG**: Oxidative, PI3K, MAPK, apoptosis, NF-k, TNF, JAK, STAT, Cytokine, Inflammation, Th17, Th1, IL-.

**Keywords REACT**: Pathways of neurodegeneration, Prion, Alzheimer, Parkinson, ALS, axon

**Keywords WIKI**: autism, thyroid hormone

These keywords were selected to specifically address a targeted hypothesis, namely the effect of inflammation on ASD (21,22,24). They best characterize the pathways and biological processes that are known to be implicated in inflammation and immunity (to the best of our knowledge).

The keywords used for searching **drug MOAs** from CMAP were:

**Keywords (MoAs)**: dopamine receptor antagonist, choline, histamine, serotonin. Once, again these keywords aim to capture the main drug MOAs that have been associated with ASD therapeutics. The full list of pathways and drug MOAs (data not shown) obtained from the search was then used as input for our methodology to perform cell-ranking.

The results from the rankings of cell types using the hypothesis-driven approach do not appear to be very much aligned with the results reported by the authors who first published the ASD dataset (2). The authors report top differentially expressed neuronal genes down-regulated for ASD in L2/3 excitatory neurons and vasoactive intestinal polypeptide (IN-VIP)–expressing interneurons. The top genes differentially expressed in non-neuronal cell types were up-regulated for ASD in protoplasmic astrocytes and microglia. They also report cell types that are recurrently affected across multiple patients as upper (L2/3) and deep layer (L5/6-CC) cortico-cortical projection neurons. This is not observed in the results of this approach whereby, Neu-NRGN-I and II cells attained the highest ranking with respect to the rest of the cell types in the analysis (see Figure 5B). Although these cells were not highlighted by the authors of the autism dataset, these results are in alignment with the CellChat rankings attained by our methodology (see Figure 4D).

### Case study 3 – COVID-19

#### Discovery Mode Approach

Although SARS-CoV-2 primarily targets the respiratory system, patients and survivors of COVID-19 can suffer neurological symptoms. A recent study performed scRNA-Seq on post-mortem brains of COVID-19 patients with the aim of investigating long-COVID (1). They revealed interesting themes linking neuroinflammation signals relayed to the brains of COVID-19 patients, particularly affecting microglia, as well as astrocytic cells. Moreover, the authors also report the neuronal subtypes that were mostly affected by these inflammatory signals. They narrow down these neuronal subtypes to gene-expression changes of the excitatory neurons, specifically L2/3 and L2/3-residing VIP interneurons. They also report microglia and astrocyte as the main cell types for relaying signals between brain barriers.

Furthermore, they show links of COVID-19 affected gene expression changes with neurological disorders such as schizophrenia, depression, Alzheimer’s disease, multiple sclerosis, Huntington’s disease and ASD.

As with the above case study, prior knowledge is included as a term list and our methodology was applied. The prior knowledge list of terms for the brain COVID-19 dataset was obtained by querying MalaCards with the keyword “Covid-19”.

The list of terms best characterizes the drugs, pathways and biological processes that are known to be implicated in COVID-19 (with respect to MalaCards). The full list of pathways and drugs (see supplementary Tables S11-15) obtained from the search was then used as input for our methodology to perform cell-ranking.

Results show that macrophages and astrocytes obtained the highest ranking according to our methodology. This is in alignment with the results obtained by the main authors of this dataset, which report that microglia and astrocytes are associated with the immune landscape of the brain in individuals with COVID-19. Furthermore, excitatory neurons received the third highest-ranking (see Figure 4E). This is also in agreement to the neuronal subtypes reported by the authors of the COVID-19dataset publication, that were narrowed down to gene-expression changes of the excitatory neurons, specifically L2/3 and L2/3-residing VIP interneurons. Enrichment analysis and DR results for some of the COVID-19 top ranked cells are shown as supplementary Figure S8-11. Results from the COVID-19 bulk RNA-Seq simulation is also shown in the box-plots (see Figure 4E), attaining an average ranking with respect to the individual cell-types in the analysis.

Results from COVID-19 and control CellChat networks, highlighted endothelial cells as the cell-types with the most cell-cell communication differences between control and disease conditions (see Figure 4F). These cell types showed a >3-fold increase in the number of normalized interactions in the COVID-19 compared to the control network (see Figure 4F). Astrocytes also appear to have cell-cell communication differences between control and disease conditions with a >1.3-fold decrease in the number of normalized interactions between networks. Again, results are in agreement to results reported by the original authors of this dataset, whereby astrocytes were proposed as one of the main cell types for relaying signals between brain barriers in these COVID-19 patient samples.

#### Hypothesis-Driven Approach

As with the above scenarios, prior knowledge is included as keywords and our methodology was applied. The prior knowledge keywords for the brain COVID-19 dataset in order to search pathway databases (KEGG, GOBP, MSIG Hallmark, REACTOME, WIKI) were:

**Keywords KEGG**: Corona disease, viral, Prion, Alzheimer, Parkinson, ALS, axon

**Keywords GO**: axon, synapse, neurotransmitter, neuron, PI3K, MAPK, apoptosis, NF-k, TNF, JAK, STAT, Cytokine, Inflammation, Th17, Th1, IL-

**Keywords MSIG**: Oxidative, PI3K, MAPK, apoptosis, NF-k, TNF, JAK, STAT, Cytokine, Inflammation, Th17, Th1, IL-.

**Keywords REACT**: SARS-CoV-2, Infectious disease, Innate Immune, Alzheimer, Huntington, autism.

**Keywords WIKI**: SARS-CoV-2, Infectious disease, Innate Immune, Alzheimer, Huntington, autism

These keywords were selected in order to address the long COVID effects with respect to inflammation in the brains of infected patients. They characterize the pathways and biological processes that are known to be implicated in inflammation as well as long COVID (to the best of our knowledge). The keywords used for searching **drug MOAs** from CMAP were:

**Keywords (MoAs)**: dopamine receptor antagonist, choline, histamine, serotonin. Once, again these keywords aim to capture the main drug MOAs that have been implicated in neurological disorders that as current research suggests, show multiple similarities to long COVID. The full list of pathways and drug MOAs (data not shown) obtained from the search was then used as input for our methodology to perform cell-ranking.

Results show that top ranked cells were epithelial and mesenchymal cells with excitatory neurons receiving the third highest-ranking (as with the Discovery mode). Keywords used were broadly associated with COVID-19 (e.g., Corona disease) as well as some targeted keywords of inflammation (e.g., Cytokine – see full list above) (see Figure 5C). These results are not in agreement with the neuronal subtypes reported by the authors of the COVID-19 dataset publication (1), which, were narrowed down to gene-expression changes of the excitatory neurons, as well as disease related astrocytes and microglia. Perhaps an explanation for these differences could be that the authors analyse the samples attained from the different brain areas (cortex and choroid plexus) separately, while we integrated all samples in our scRNA-Seq analysis.

### Comparing to standard cell type ranking methods (proportion of cells and number of differentially expressed genes)

Standard cell ranking methods, such as comparison of the total number of DEGs between disease/control conditions for individual cell types, are often used to prioritize cell types and highlight cells which can be deemed important for a specific disease (2). The calculation of the proportion of cell types across the different conditions in a a scRNA-Seq experiment, is yet another standard way of cell type prioritization for a targeted disease. In order to a assess the differences between these standard approaches and our methodology we proceeded to generate results using these methods for all three datasets used as case studies and compared them to our results.

Comparing results of our ranking methodology to the ranking generated from standard methods used for scRNA-Seq ranking, we observe certain similarities but also some discrepancies. For example, for the LAM dataset it appears that smooth muscle cells are not ranked high with respect to the proportional differences between disease donor conditions (see Figure 6A). Perhaps the number of cells is not affected in an analogous manner to the molecular perturbations which are commonly seen in LAM disease (17–19). Similarly, DEG analysis for these cells does not show major differences in the number of DEGs between disease and donor conditions (see Figure 7A). For the AUTISM dataset the top ranked cell types with respect to proportional differences between disease and control conditions are oligodendrocytes while the cells which are reported in the main primary work that published this dataset (i.e., L2/3, IN-VIP) are found lower down in the rankings (see Figure 6B). The same pattern is observed in the number of DEGs between disease and control samples for this dataset (see Figure 7B). For the COVID-19 brain dataset, the top ranked cell types were epithelial and neural cells for the proportion of cells and DEGs analyses respectively (see Figure 6C and 7C). The macrophage and astrocytes cells reported in the main research paper for this dataset appear lower down in the rankings. While the excitatory neurons deemed as highly informative by the authors of this dataset appear high in the rankings of the proportional cell analysis but lower in the rankings of the DEGs analysis (see Figure 6C and 7C).

**Figure 6.**
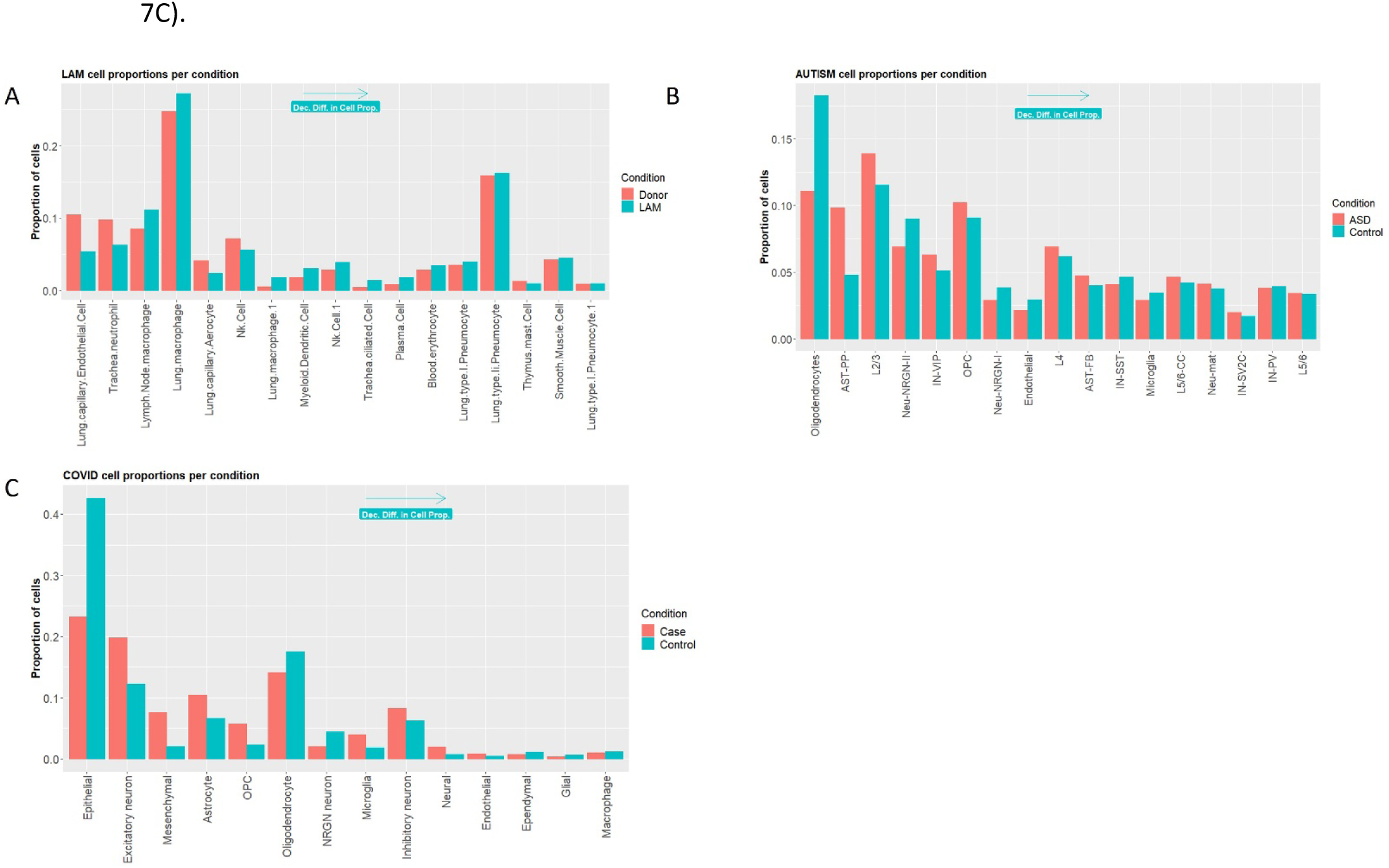
Proportion of cell types across different conditions for the 3 datasets used to validate our methodology. **A.** LAM disease dataset and the proportion of cell types across conditions. **B.** AUTISM disorder dataset showing the proportion of cell types across conditions. **C.** Brain COVID-19 dataset showing the proportion of cell types across conditions. Cells are ranked in descending order with respect to the difference between the proportion of cells in disease vs control samples.

**Figure 7.**
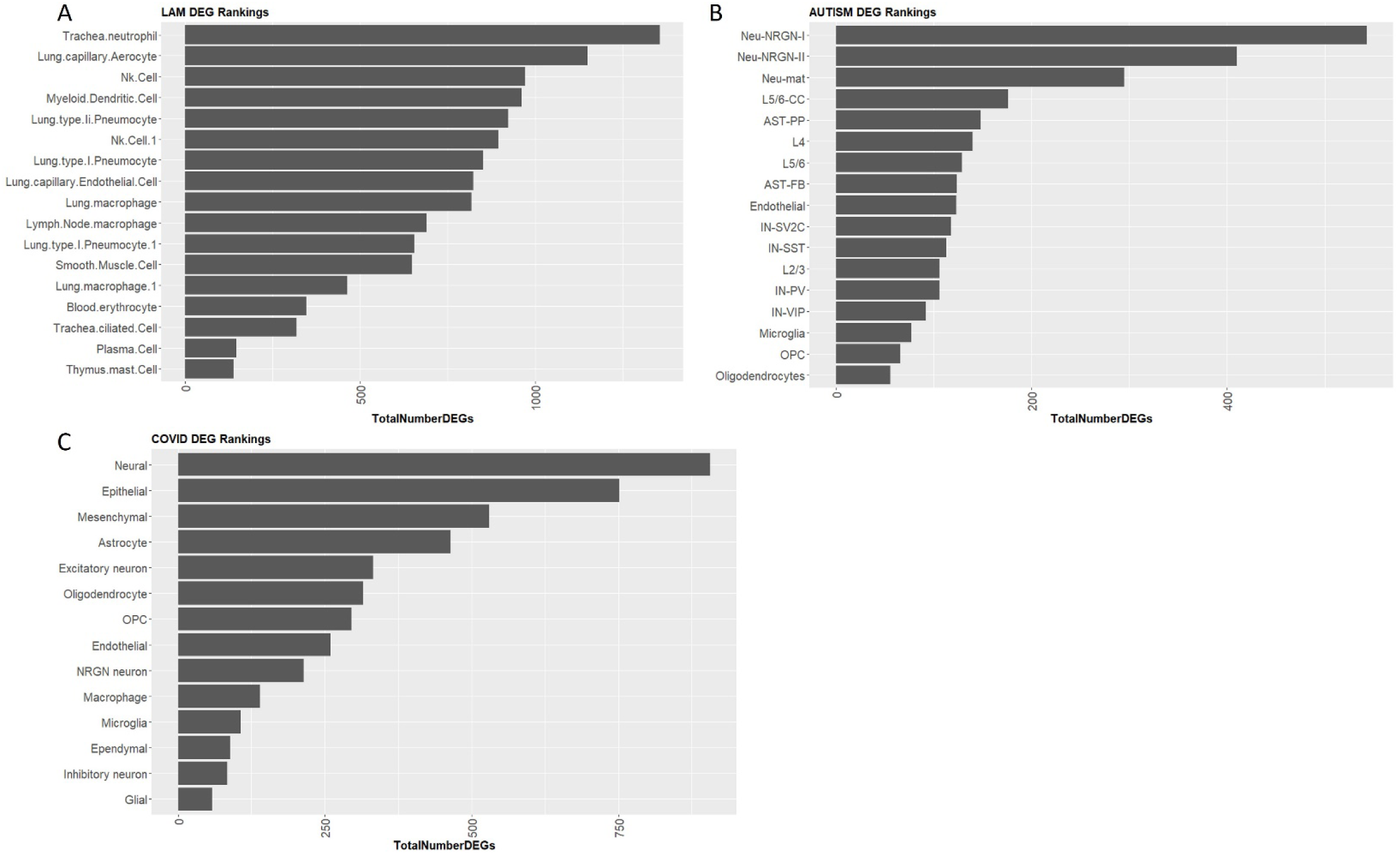
Number of differentially expressed genes (DEGs) for each cell type across different conditions for the 3 datasets used to validate our methodology. **A.** LAM disease dataset and the number of DEGs for individual cell types across conditions. **B.** AUTISM disorder dataset showing the number of DEGs for individual cell types across conditions. **C.** Brain COVID-19 dataset showing the number of DEGs for individual cell types across conditions. Cells are ranked in descending order with respect to the total number DEGs in control vs disease samples.

## Discussion

Currently the use of methods such as, total number of differentially expressed genes (DEGs) between disease/control conditions for individual cell types, are utilized in order to obtain an indication as to which cell types are more profoundly implicated in the disease under study (2). These methods have certain drawbacks and limitations as shown by assessments in both simulated and experimental datasets. Specifically, the number of DEGs is often strongly correlated with the number of cells per type, causing abundant cell types with modest transcriptional perturbations to be prioritized over rare but more strongly perturbed cell types (4). Similarly, counting the number of different cell types between disease/control conditions and calculating the proportion of cells between conditions can also be an indication of the cellular differentiation in disease vs. control. For example, in a recent study by *Yang et al* (1) the authors use differences in proportion of cells between macrophages in disease *vs.* control conditions to pin-point specific disease associated macrophages. This again, can be a very important indication as to which cell types are mostly affected by the disease. However, this method relies heavily on accurate normalization procedures to ensure that all libraries are contributing equally towards this assessment criterion. Also cell type proportion estimates from scRNA-seq data are variable, and statistical methods that can correctly account for different sources of variability are needed to confidently identify statistically significant shifts in cell type composition between experimental conditions (4). Moreover, it may be difficult to decide on that exact number of cells that can reliably be considered as a significant difference between disease and control samples and further play a role in disease specific, causative, cellular differentiation.

We propose an approach which uses prior knowledge in an automated, structured manner. Our approach combines both prior knowledge available via disease related databases, but also allows for intervention by users which are experts for the disease under study. This permits a more targeted hypothesis-driven approach by supplying keywords by which to search available database resources. The supplied prior knowledge is further mapped to enrichment and drug repurposing results obtained from a scRNA-Seq data-driven analysis and a ranking of the individual cell clusters is performed based on the accuracy of this mapping. We perform three case study validations using published scRNA-Seq datasets and compare the results of the rankings obtained by our methodology with the results reported in the published research papers for the individual datasets. We find that our methodology is able to highlight specific cell types that are known to be implicated in the diseases under study as reported by clinical information as well as previous published results (1–3). We show that using our method can provide an automated, structured analysis pipeline that can allude to very similar results attained by relevant research which utilized more traditional, manual, time-consuming methods that require extensive and thorough investigation into the molecular basis of the disease under study. Furthermore, our methodology allows for more targeted hypothesis-driven approaches to highlight specific cell types that are implicated in molecular mechanisms relevant to the hypothesis under consideration. We show that applying specific themes (e.g., inflammation) in our hypothesis driven approach, alters cell ranking results and allows for the identification of cell types which potentially contribute towards these processes.

Overall, our methodology has substantial advantages to more traditional cell ranking techniques. However, we cannot replace other prioritization methods or exclude their potential usefulness. We provide a complementary methodology that utilizes prior knowledge in a rapid and automated manner, that has previously not been attempted by other studies. As all novel methods, it is not without limitations and specifically the non-completeness of the databases from where we are retrieving the input knowledge and the lack of uniform nomenclature can be significant limiting factors in our analysis. Moreover, the lack of agreement between prior knowledge databases and results from scRNA-Seq analysis (mapping), can also act as a limitation. The use of the hypothesis driven approach can potentially mitigate this problem, as the user can manual add or replace keywords in the prior knowledge checklist. This can allow for better mapping between terms derived from prior knowledge with those from data-driven results.

Our methodology is available as an R package (scRANK) that is freely available for download and installation (https://github.com/aoulas/scRANK).

## Author Contributions

AO and GMS designed the study. AO, NK and KS conceived and developed the computational tools. AO analyzed the data and interpreted the results. NK, KS and GMS interpreted the results. KS contributed to the drug database data for the analysis. AO designed the experiments and wrote the manuscript. AO and GMS conceived the project. All authors edited the manuscript.

